# Transcriptional activation of cyclin D1 via HER2/HER3 contributes to cell survival and EGFR tyrosine kinase inhibitor resistance in non-small cell lung carcinoma

**DOI:** 10.1101/851279

**Authors:** Bin Liu, Deng Chen, Shipeng Chen, Ali Saber, Hidde Haisma

## Abstract

Several different mechanisms are implicated in the resistance of lung cancer cells to epidermal growth factor receptor-tyrosine kinase inhibitors (EGFR-TKIs), and only few have been functionally investigated. Here, using genetically knocked out *EGFR* and TKI-resistant lung cancer cells, we show that loss of wild-type *EGFR* attenuates cell proliferation, migration and 3D-spherid formation, whereas loss of mutant *EGFR* or resistance to TKIs reinforces those processes. Consistently, disruption of wild-type *EGFR* leads to suppression of HER2/HER3, while mutant *EGFR* ablation or resistance to TKIs increases HER2/HER3 expression, compensating for EGFR loss. Furthermore, HER2/HER3 nuclear translocation mediates overexpression of cyclin D1, leading to tumor cell survival and drug resistance. Cyclin D1/CDK4/6 inhibition resensitizes erlotinib-resistant (ER) cells to erlotinib. Analysis of cyclin D1 expression in patients with non-small cell lung carcinoma (NSCLC) showed that its expression is negatively associated with overall survival and disease-free survival. Our results provide biological and mechanistic insights into targeting EGFR and TKI resistance.

## Introduction

Lung cancer is the leading cause of cancer mortality worldwide. In 2018, more than 2 million new cases and 1.7 million lung cancer-related deaths were reported^1^. Non-small cell lung carcinoma (NSCLC) is the main type of lung cancer, accounting for 85% of all lung cancer cases^2^. Surgery or surgery followed by chemotherapy is the common choice for patients with resectable lung cancer. For those with advanced stage cancer, targeted therapy can be used to achieve a longer progression-free survival^2^.

Tyrosine kinase inhibitors (TKIs), such as erlotinib, gefitinib, and afatinib, are the commonly used targeted agents for NSCLC patients with certain *EGFR* mutations^3, 4^. EGFR-TKIs bind to the tyrosine kinase domain of EGFR, blocking tyrosine kinase activity and downstream signaling. Approximately 10% of NSCLC patients from Western countries and 30% of NSCLC patients from East Asia, harboring *EGFR*-activating mutations (exon 19 deletions and L858R point mutation), could benefit from TKIs treatment^5, 6^. However, inevitably, almost all patients treated with TKIs develop resistance within 9-13 months after treatment initiation^3, 7, 8^.

Mechanisms of EGFR-TKI resistance in NSCLC are complicated. The gatekeeper mutation T790M contributes to almost 50% of all acquired resistance to elortinib and gefitinib. Activation of compensatory signaling pathways, such as c-MET, PI3K/AKT/mTOR, JAK2/STAT3, are other mechanisms of resistance against TKIs^3, 9, 10^. Furthermore, dysregulation of HER family receptors and EGFR nuclear trans-location are reported as mechanisms of resistance^11^. EGFR nuclear translocation is considered a transcriptional co-activator of several well-known transcription factors, such as STAT3/5^12, 13^, E2F1^14^ and RNA helicase A^15^, regulating the expression of many essential genes, including cyclin D1. Although the function of nuclear EGFR is relatively clear, functions of other nuclear HER family receptors, including HER2/3/4, remain unknown in NSCLC.

In this study, we aimed to characterize cellular and molecular changes of HER family receptors upon CRISPR/Cas9-mediated disruption of *EGFR* or acquisition of resistance to EGFR-TKIs. We showed that wild-type *EGFR* ablation results in a substantial inhibition of cell proliferation and migration, but that disruption of mutant *EGFR* or acquisition of TKI resistance promoted cell proliferation and migration resulting in cell survival. Furthermore, we found that upregulation of HER2 and HER3 mediated cyclin D1 overexpression, contributing to cell survival and proliferation upon EGFR loss or emergence of TKI-resistance.

## Results

### Establishment of *EGFR^-/-^* and EGFR-TKI resistant cell lines

To conduct clinically relevant studies of EGFR-positive lung cancer cells, we established *EGFR****^-/-^*** and EGFR-TKI resistant cell lines. Four different NSCLC cell lines (A549, H1299, H1650 and HCC827) with different genotypes were included in this study (Table 1). The A549, H1299, H1650 *EGFR****^-/-^*** cell lines were generated by using CRISPR/Cas9 gene editing technology (Supplementary Fig. 1A)^16, 24^.

**Table 1.** List of NSCLC cell lines used in this study.

| **Cell line** | **Lung cancer subtype** | ***EGFR* status** |
| --- | --- | --- |
| A549 | Adenocarcinoma | Wild-type |
| H1299 | Adenocarcinoma | Wild-type |
| H1650 | Adenocarcinoma | Exon19 del |
| HCC827 | Adenocarcinoma | Exon19 del |

A549 and H1299 contain wild-type *EGFR* and are resistant to the erlotinib (an EGFR-TKI), while H1650 and HCC827 harbor an *EGFR* exon 19 deletion, which is the most common *EGFR* mutation and makes tumor cells sensitive to TKIs. We established H1650-ER and HCC827-ER cell lines after long-term exposure to erlotinib (Supplementary Fig. 1B and 1C). For HCC827ER, IC_50_ increased dramatically from 0.1 ±0.18 μM to 47.6 ±0.17 μM. H1650ER cells were able to grow in high dose (50 μM) of erlotinib without loss of viability, although the parental cells had moderate response to erlotinib. Furthermore, we observed an aggressive pattern of hyper-progression when H1650ER cells were treated with erlotinib (Supplementary Fig. 1C).

**Fig. 1.**
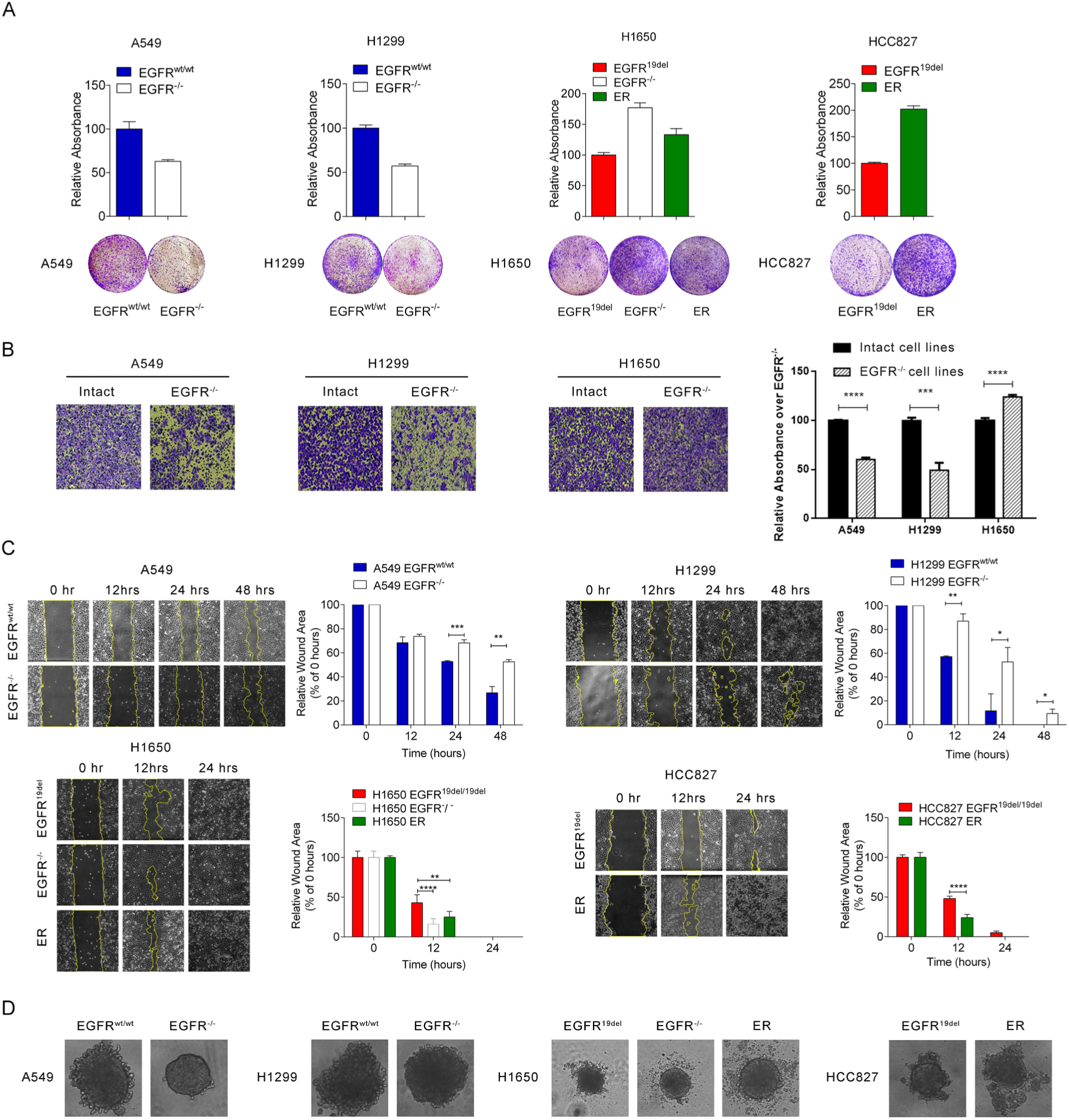
Characterization of EGFR loss and erlotinib resistant phenotypes in different NSCLC cell lines. **A.** Cell proliferation determined by colony formation assay. **B.** Transwell assay to measure vertical migration. **C.** Wound healing assay to measure lateral migration. **D.** 3D-spheroid tumor formation with U-bottom ultra-low attachment culture.

### Wild-type *EGFR* ablation attenuates cell proliferation, migration and 3D spheroid formation, but disruption of *EGFR^19del^* or acquired resistance accelerates them

To elucidate the effect of *EGFR* ablation and TKI-resistance on proliferation and migration of lung cancer cells with different *EGFR* mutational statuses, we performed colony formation, trans-well, wound healing and 3D spheroid tumor formation assays.

In *EGFR^wt/wt^* cell lines (A549 and H1299), *EGFR* ablation significantly inhibited cell proliferation, vertical and horizontal migration and tumor formation. Cell proliferation of *EGFR-*KO cells decreased by 50% compared to the parental cells (Fig. 1A). Similarly, impaired ability of migration was observed in trans-well and wound healing assays (Fig. 1B and 1C). In 3D-spheroid tumor formation assay, original A549 and H1299 cells formed larger tumors and showed stronger invasion ability than corresponding *EGFR^-/-^* cells (Fig. 1D). In contrast, we observed opposite results in *EGFR^19del^* and erlotinib-resistant cells. In H1650 and HCC827, *EGFR^19del^* ablation or acquired erlotinib resistance led to acceleration of cell proliferation, vertical and horizontal migration, and 3D-spherid formation (Fig. 1A-D).

### Distinct changes in pERK(1/2) and pAKT levels upon *EGFR* knockout and acquired TKI resistance

*EGFR* status can significantly affect two main downstream pathways, i.e. the MAPK/pERK(1/2) and PI3K/AKT. *EGFR*-activating mutations, overexpression or inhibition can dramatically affect these two pathways and substantially influence cancer cell growth and survival. To precisely investigate the effect of different *EGFR* status on these two pathways, we performed a panel of western blot to detect changes in the levels of phosphorylated ERK (pERK)(1/2) and phosphorylated AKT (pAKT) in different NSCLC cell lines.

We found significant changes in pERK(1/2) and pAKT levels upon *EGFR* ablation or TKI treatment. pAKT expression level was higher in A549 *EGFR^-/-^* (Fig. 2A), but downregulated in H1299 *EGFR^-/-^* as compared to parental cells (Fig. 2B). Interestingly, A549 *EGFR^-/-^* showed significant increases in both AKT and pAKT levels in comparison with A549 *EGFR^wt/wt^*. However, the increased levels of pAKT in A549 *EGFR^-/-^* were reduced upon elortinib treatment (Fig. 2A). Strikingly, pERK levels significantly increased when we treated A549 *EGFR^-/-^* with elortinib (Fig. 2A), which was not observed in any other cell lines (Fig. 2B, 2C and 2D). Similar to A549, pAKT levels were upregulated in H1650 *EGFR^-/-^* and HCC827ER, but elortinib treatment restrained pAKT levels (Fig. 2C and 2D).

**Fig. 2.**
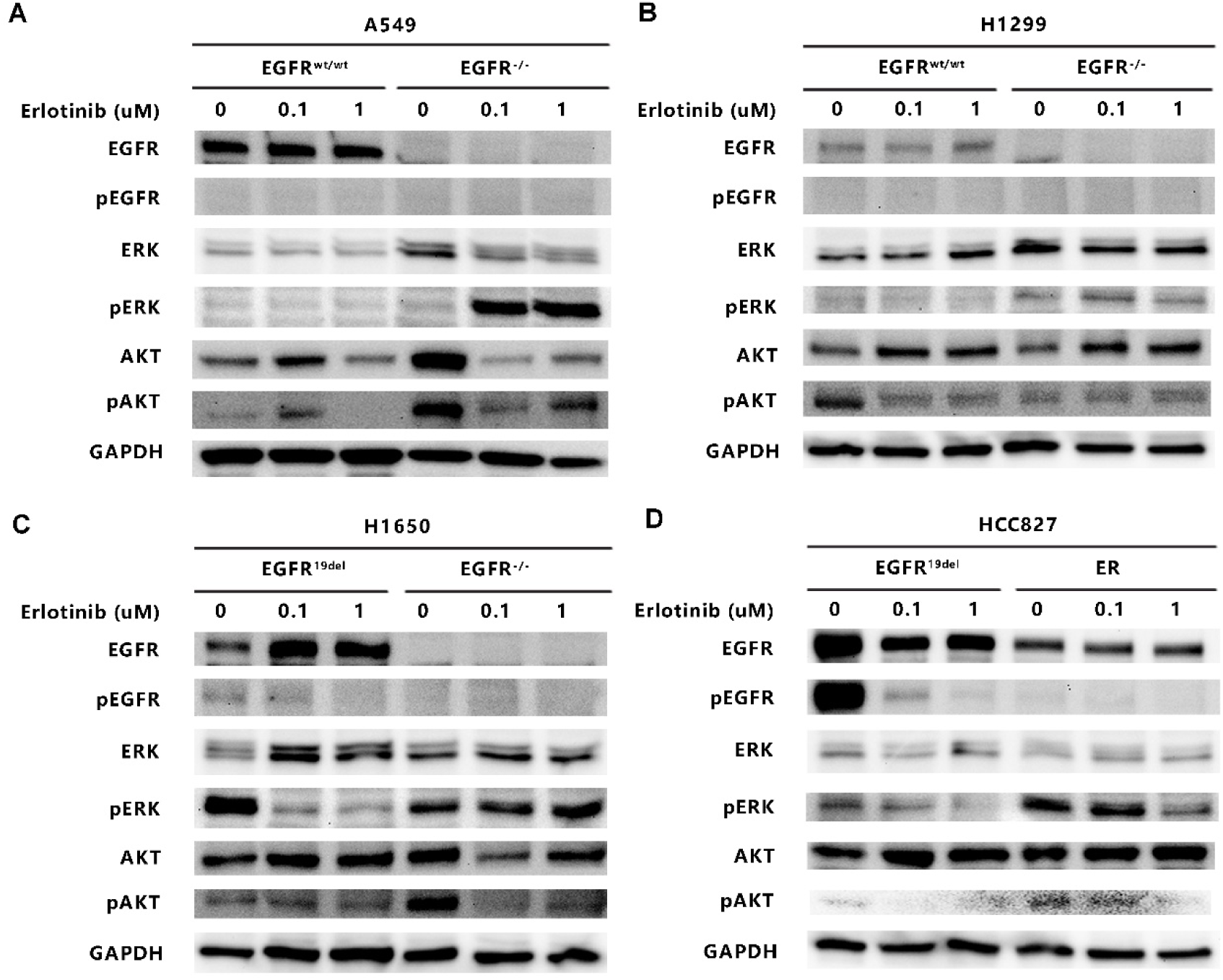
Western blot analysis of downstream pathways after elortinib treatment in *EGFR* knockout, erlotinib resistant and original NSCLC cells. **A.** Western blot analysis of A549 *EGFR^wt/wt^* and *EGFR^-/-^* cells. **B.** Western blot analysis of H1299 *EGFR^wt/wt^* and *EGFR^-/-^* cells. **C.** Western blot analysis of H1650 *EGFR^del^* and *EGFR^-/-^* cells. **D.** Western blot analysis of HCC827 and HCC827ER cells.

### HER2 and HER3 are upregulated in *EGFR^19del^* ablated and/or elortinib resistant cells, but not in *EGFR^wt/wt^* disrupted cells

Dysregulation of HER family receptors can occur in patients with NSCLC upon TKIs treatment^11^. Precise evaluation of HER family receptor expression levels upon and during treatment with TKIs can provide deeper understanding of mechanisms of TKI resistance. Therefore, we investigated alterations in HER2 and HER3 expression levels upon *EGFR* ablation and EGFR-TKI resistance.

In *EGFR*^wt/wt^ cell lines (A549 and H1299), *EGFR* disruption resulted in a slight decrease in HER2 and HER3 levels (Fig. 3A and 3B). However, cells with mutant *EGFR*, i.e. H1650 and HCC827, showed a different pattern. HER2 and HER3 were dramatically upregulated in H1650 *EGFR^-/-^* and H1650-ER cells at both mRNA and protein level compared to the parental cells (Fig. 3A and 3B). In HCC827-ER cells, only HER2 protein level was upregulated (Fig. 3B). Although HER3 was significantly upregulated in HCC827-ER at RNA level, this was not reflected at protein level, suggesting that post-transcriptional regulation mechanisms may be involved.

**Fig. 3.**
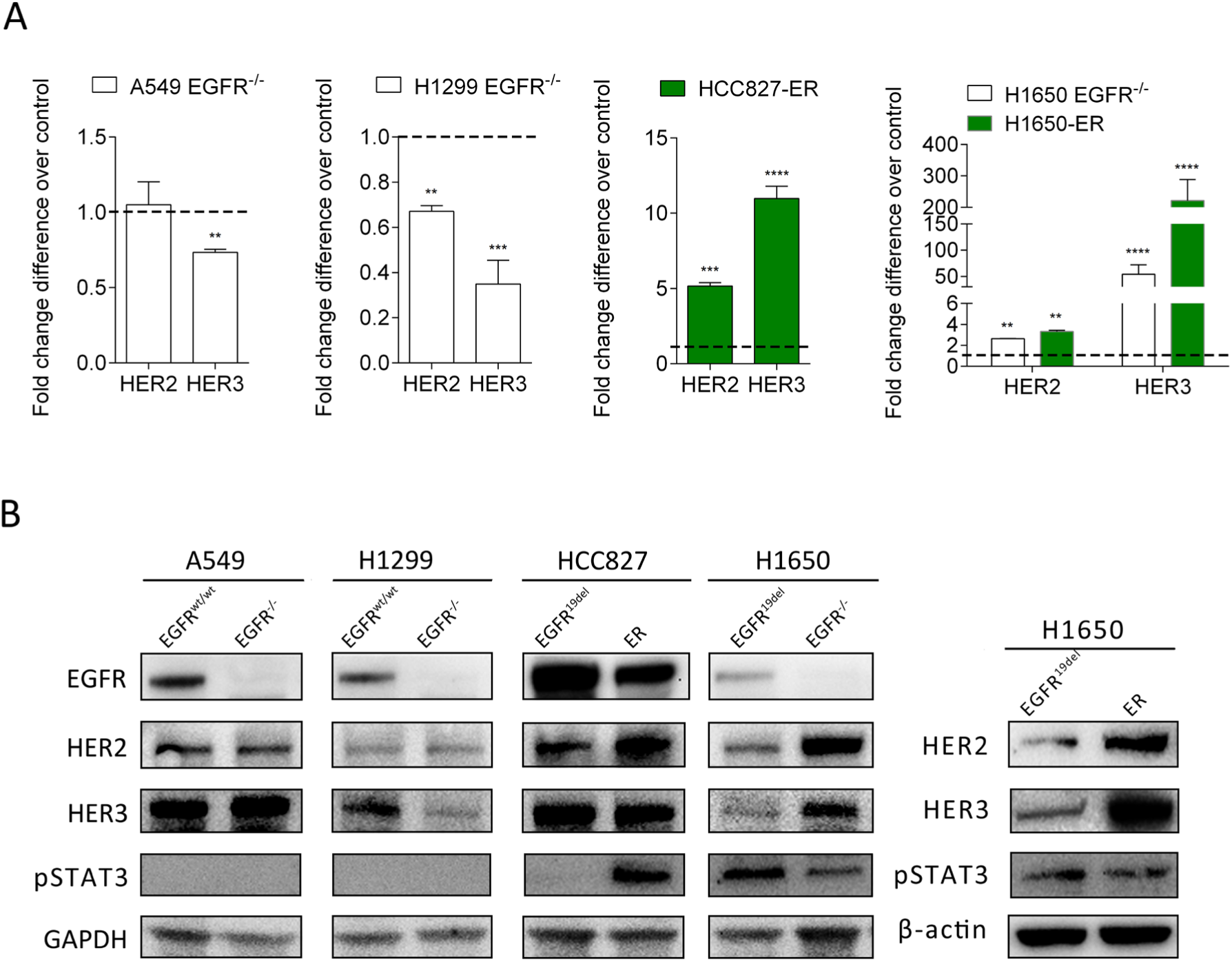
Expression levels of HER2 and HER3. **A.** Relative mRNA expression of HER2 and HER3 as compared to parental cells. **B.** Western blot analysis of EGFR, HER2, HER3 and pSTAT3 in different NSCLC cell lines.

In addition, we evaluated pSTAT3 levels, since it is an important downstream protein of HER family receptors. In *EGFR^wt/wt^* cell lines, pSTAT3 was undetectable (Fig. 3B), while pSTAT3 level was significantly increased in HCC827ER compared to HCC827 cells (Fig. 3B). Together, these results suggest that upregulation of HER2 and HER3 occurs only in mutant *EGFR* knockout and TKI-resistant NSCLC cell lines, but not in wild-type *EGFR* knockout cells.

### Pan-HER family inhibitor, afatinib, suppresses pAKT via inhibition of HER2 and HER3

HER2 and HER2-containing heterodimers have the strongest kinase activity compared to other HER family receptors, which makes them an important therapeutic target in patients with cancer. Afatinib not only can block EGFR, but also inhibits the tyrosine kinase activity of HER2 and other HER family receptors. Since we showed that HER2 and HER3 were upregulated in *EGFR^19del^* disrupted and ER cells, we speculated that suppression of HER2 and HER3 may further inhibit tyrosine kinase activity and tumor proliferation. Therefore, we treated H1650 *EGFR^-/-^* and HCC827ER cells with afatinib to block HER2- and HER3-mediated signaling. We observed a decreased level of pAKT upon afatinib treatment in H1650 *EGFR^-/-^* and HCC827-ER cells (Fig. 4A and 4B). However, we did not observe significant inhibition of cell proliferation, although the downstream pAKT signaling was decreased by afatinib. Thus, these results suggest that overexpression of HER2 and HER3 may play other functions than tyrosine kinase activity in H1650 *EGFR^-/-^* and HCC827-ER cells.

**Fig. 4.**
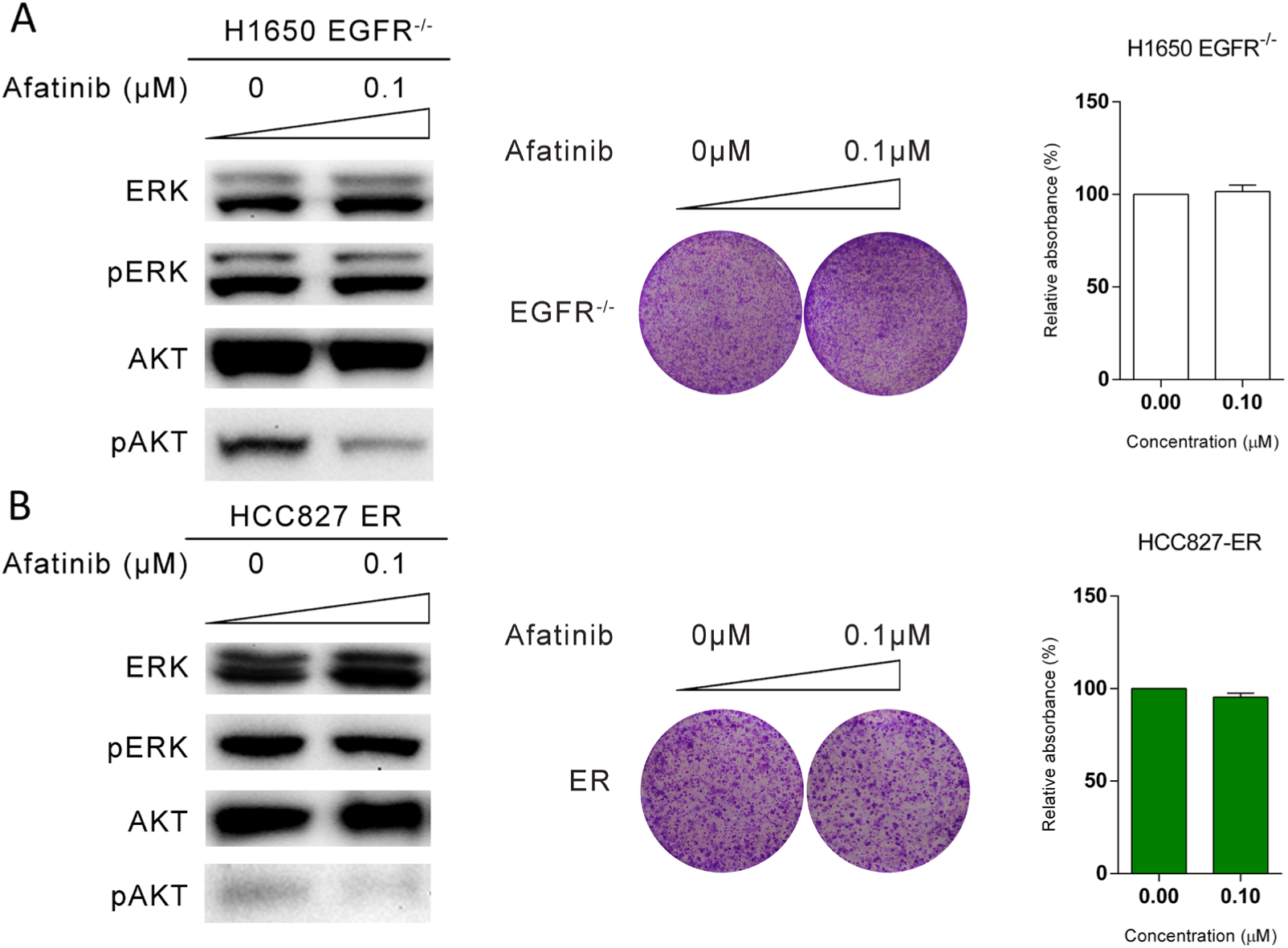
Effects of afatinib treatment on MAPK/ERK and PI3K/AKT signaling and the cell proliferation. **A.** MAPK/ERK and PI3K/AKT signaling and cell proliferation with treatment of afatinib in H1650 and H1650 EGFR^-/-^. **B.** MAPK/ERK and PI3K/AKT signaling and cell proliferation with treatment of afatinib in HCC827 and HCC827 ER.

### HER2 and HER3 are overexpressed in the nuclei of *EGFR^19del^* ablated and ER cells

EGFR nuclear translocation is one of the mechanisms of resistance to TKI in NSCLC. Whether nuclear translocation of other HER family receptors is also involved in EGFR-TKIs resistance is unclear. Hence, we assessed HER2 and HER3 expression levels in non-nuclear and nuclear fraction in our NSCLC cell line panel. We found that HER2 and HER3 were mainly expressed in the nuclei in all tested NSCLC cell lines (Fig. 5A-E). Ablation of *EGFR^wt/wt^* influenced neither HER2 and HER3 expression, nor their nuclear-membrane distribution in A549 cells (Fig. 5A). Similarly, we did not observe significant changes of HER2 and HER3 in H1299 *EGFR^wt/wt^* and *EGFR^-/-^* cells due to the relatively low expression of protein (Fig. 5A). Interestingly, HER2 and HER3 were both significantly overexpressed in the nuclei of *EGFR^19del^* disrupted and ER cells compared to the parental cells H1650 and HCC827 (Fig. 5C-E).

**Fig. 5.**
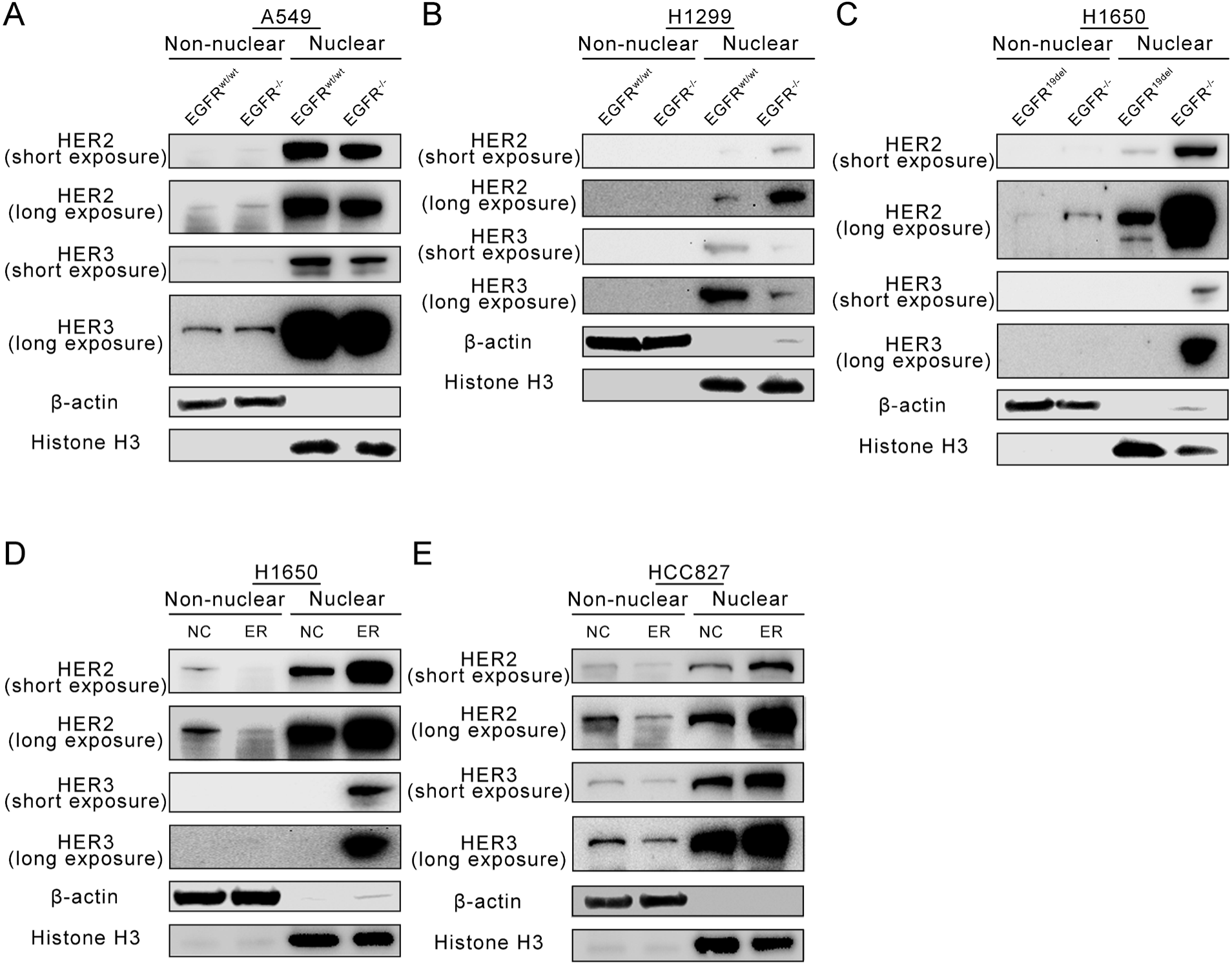
Western blot analysis of HER2 and HER3 expression levels in non-nuclear and nuclear fractions of NSCLC cells. **A.** Western blot analysis of HER2 and HER3 expression levels in non-nuclear and nuclear fractions from A549 and A549 *EGFR^-/-^* cells **B.** Western blot analysis of HER2 and HER3 expression levels in non-nuclear and nuclear fractions from H1299 and H1299 *EGFR^-/-^* cells. **C.** Western blot analysis of HER2 and HER3 expression levels in non-nuclear and nuclear fractions from H1650 and H1650 *EGFR^-/-^* cells. **D.** Western blot analysis of HER2 and HER3 expression levels in non-nuclear and nuclear fractions from H1650 and H1650 ER cells. **E.** Western blot analysis of HER2 and HER3 expression levels in non-nuclear and nuclear fractions from HCC827 and HCC827 ER cells.

### Transcriptional complex of HER2/HER3/pSTAT3 increases in nuclei, enriches in the cyclin D1 promoter region and reinforces cyclin D1 expression in *EGFR^19del^* ablated and ER cell lines

HER2/HER3/Stat3 can form a transcriptional complex and stimulate cyclin D1 expression in breast cancer^20, 21^. However, this mechanism has not yet been confirmed in lung cancer. Therefore, we performed co-immunoprecipitation assays to explore this mechanism. Our results showed that in *EGFR^wt/wt^* cell lines, *EGFR* deletion did not influence HER2/HER3/pSTAT3 heterotrimer formation in the nucleus (Fig. 6A and 6B). However, in *EGFR*-mutant cell lines, *EGFR* ablation or ER led to HER2/HER3/pSTAT3 nuclear enrichment (Fig. 6C and 6D).

**Fig. 6.**
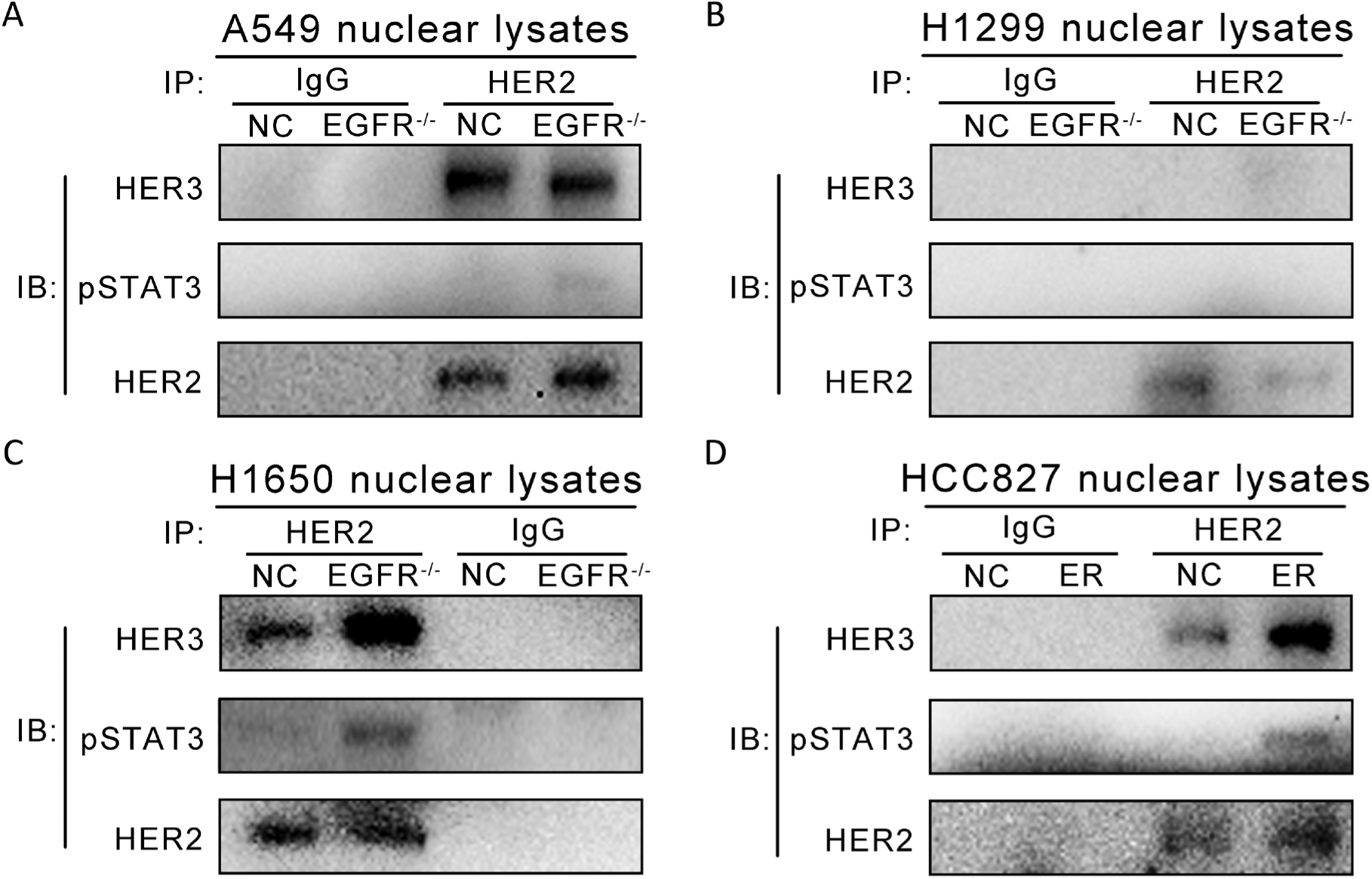
Co-Immunoprecipitation (Co-IP) analysis of HER2/3/pSTAT3 heterotrimer in NSCLC nuclei. **A.** Co-Immunoprecipitation analysis of HER2/3/pSTAT3 heterotrimer from A549 and A549 *EGFR^-/-^* nuclear lysates. **B.** Co-Immunoprecipitation analysis of HER2/3/pSTAT3 heterotrimer from H1299 and H1299 *EGFR^-/-^* nuclear lysates. **C.** Co-Immunoprecipitation analysis of HER2/3/pSTAT3 heterotrimer from H1650 and H1650 *EGFR^-/-^* nuclear lysates. **D.** Co-Immunoprecipitation analysis of HER2/3/pSTAT3 heterotrimer from HCC827 and HCC827ER nuclear lysates.

In the next step, we performed sequential ChIP assay to detect the interactions between HER2/HER3/pSTAT3 and the cyclin D1 (CCND1) promoter. We found that HER2/HER3 complex bound more to the *CCND1* promoter in HCC827ER and H1650 *EGFR****^-/-^*** cells than in original cell lines (Fig. 7A). Conversely, less binding was observed in *EGFR^wt/wt^* ablated cells than in parental cell lines (Fig. 7A). Consequently, higher cyclin D1 mRNA levels were observed in *EGFR^19del^* disrupted and ER cells compared to both parental and wild-type EGFR cells (Fig. 7B). Furthermore, western blotting showed that *EGFR^wt/wt^* ablation downregulated cyclin D1, while disruption of *EGFR^19del^* or erlotinib resistance increased its expression (Fig. 7C). Collectively, deletion of *EGFR* reduced affinity of HER2/HER3 complex to cyclin D1 promoter, decreased cyclin D1 expression in EGFR^wt/wt^ cells, but reinforced it in cells with mutant EGFR.

**Fig. 7.**
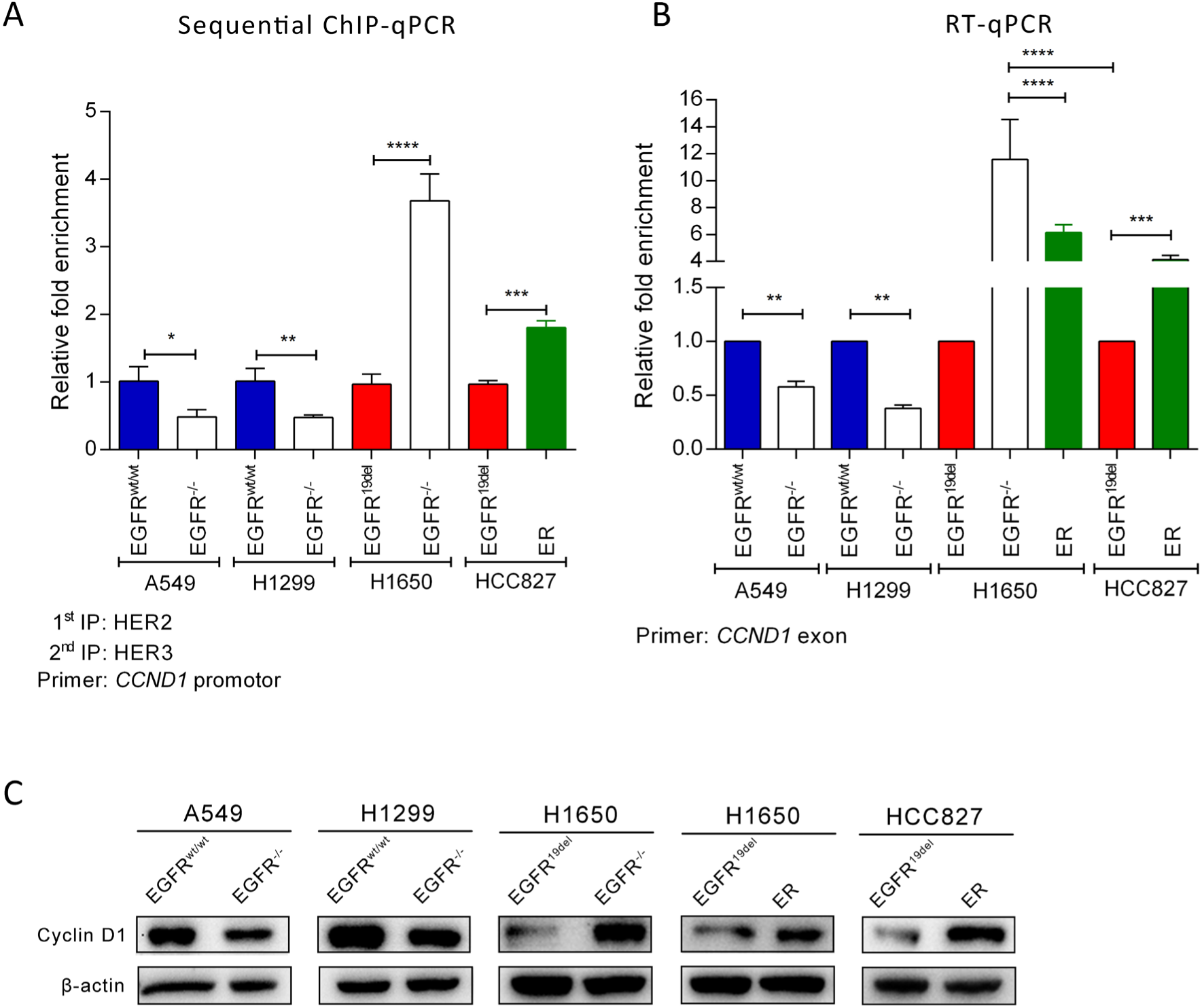
HER2 and HER3 complex binding to cyclin D1 promoter and cyclin D1 expression levels upon EGFR loss or erlotinib resistance in different NSCLC cell lines. **A.** Enrichments of HER2 and HER3 complex binding to cyclin D1 promoter detected by sequential ChIP and qRT-PCR. Chromatins were first immunoprecipitated with HER2 antibody and were re-immunoprecipitated using HER3 antibody. Primer set for sequential ChIP was designed to amplify the fragment of cyclin D1 promoter including the binding site of HER2/3/pSTAT3 complex. **B.** Relative mRNA expression of cyclin D1. **C.** Western blot analysis of cyclin D1 expression in different NSCLC cell lines.

### Ribociclib resensitizes cells to erlotinib

Although no drug is available to directly and specifically inhibit cyclin D1, an inhibitor of cyclin D1/CDK4/6 (ribociclib) has been approved by FDA in March 2017 for breast cancer treatment. We asked whether ribociclib can resensitize ER cells to erlotinib treatment. We treated *EGFR^19del^* cells (H1650ER and HCC827ER) with either erlotinib or ribociclib or combinations thereof. We observed a significant combinatorial effect of ribociclib and elortinib in H1650ER and HCC827ER cells colony formation (Fig. 8A and 8B).

**Fig. 8.**
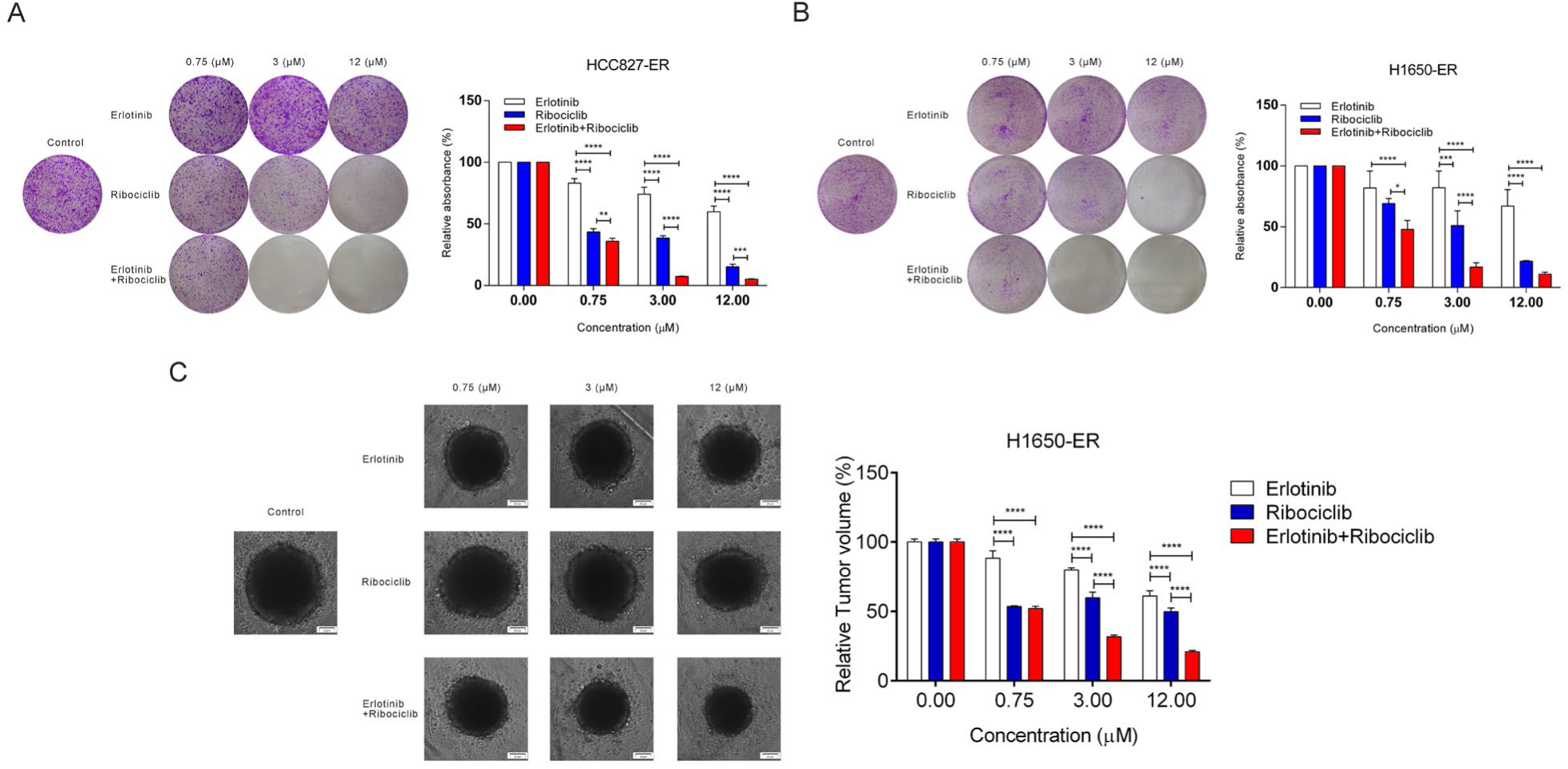
Effects of the treatment with Elortinib and/or Ribociclib in H1650ER and HCC827ER cells. **A.** erlotinib or ribociclib or the combinations thereof for HCC827ER. **B.** erlotinib or ribociclib or the combinations thereof for H1650ER. **D.** erlotinib or ribociclib or the combinations thereof for H1650ER-3D spherid tumor.

To mimic *in vivo* tumor growth, we further treated H1650ER and HCC827ER cells using a 3D spheroid model. Consistently, a significant combinatorial effect of ribociclib and elortinib was observed in H1650ER 3D-spheroid tumor (Fig. 8C), whereas HCC827ER cells are not able to form 3D spheroids with treatment.

### High expression of CCND1 is correlated with lower overall survival and disease-free survival rates in patients with lung cancer

To validate the clinical relevance of our results, we analyzed gene expression data from patients with lung cancer retrieved from The Cancer Genome Atlas (TCGA) database. Based on Kaplan-Meier survival analysis, NSCLC patients with higher cyclin D1 had a significantly lower overall survival (OS) and disease-free survival rates (DFS) (Fig. 9A). The median DFS and median OS were significantly shorter in patients with higher CCND1 expression than those with lower gene expression (2.58 vs 3.55 years, *p*=0.013; 3.52 vs 5.16 years, *p*=0.038).

**Fig. 9.**
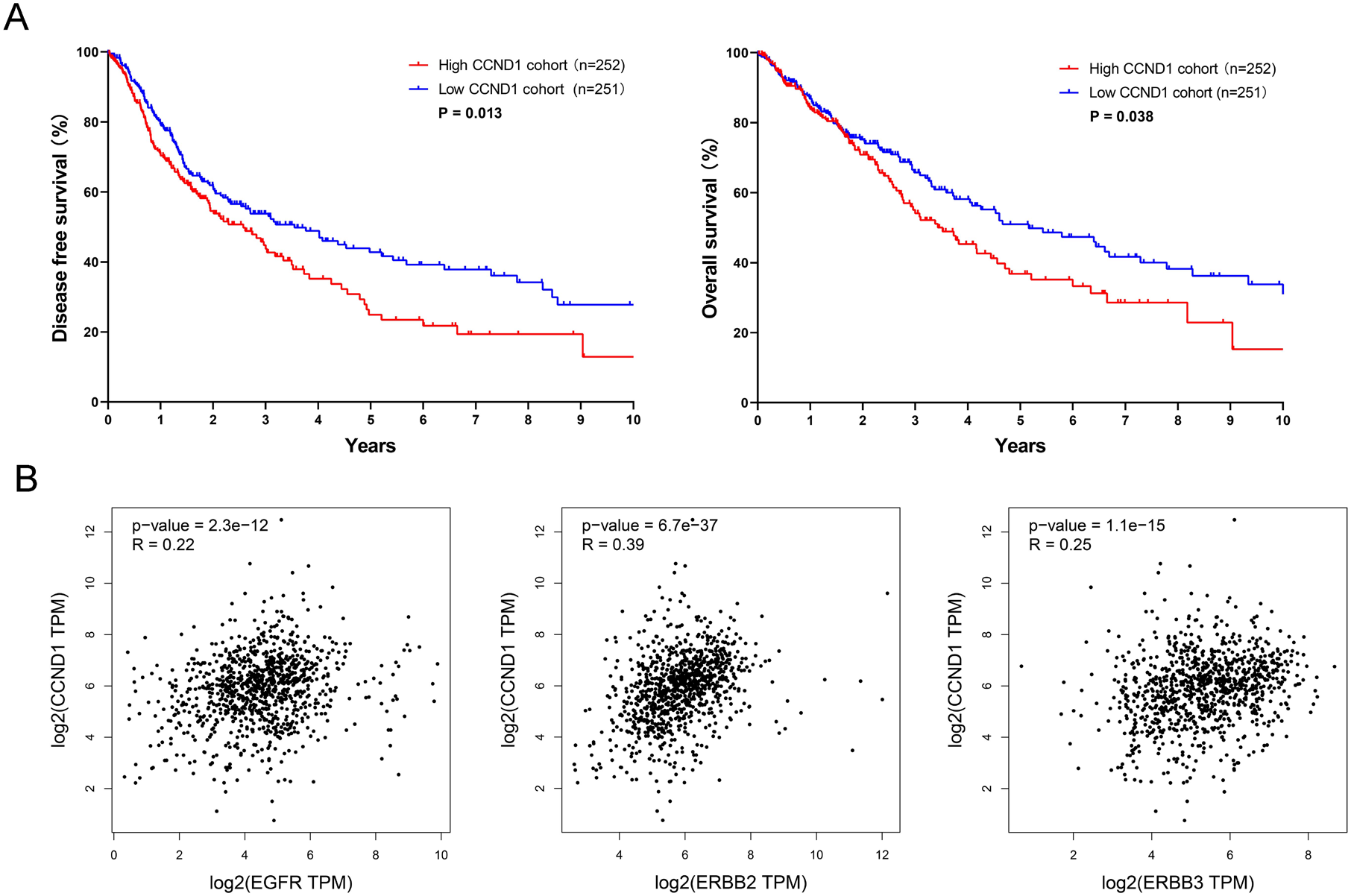
Gene expression data analysis from patients with lung cancer. A. CCND1 high expression is associated with a poor prognosis of lung cancer patients. Kaplan–Meier survival curve of disease-free survival (DFS) and overall survival (OS) in NSCLC patients with high CCND1 expression and low CCND1 expression. B. Correlation between CCND1 and Her family receptors. The correlation between CCND1 and Her family receptors based on TCGA data in GEPIA. TPM: Transcripts Per Kilobase of exon model per Million mapped reads.

Furthermore, there was a significant positive correlation between HER2 and cyclin D1 expression (Fig. 9 B), which was stronger than EGFR/ cyclin D1 or HER3/cyclin D1 correlation.

Taken together, these results suggest that cyclin D1 may be a promising therapeutic target for patients with NSCLC.

## Discussion

Epidermal growth factor receptor is a key therapeutic target for patients with Non-Small Cell Lung Cancer. Although the majority of patients with NSCLC harboring EGFR activating mutations benefit from EGFR-TKI treatment, acquired resistance inevitably arises. Extensive efforts have led us to the identification of several resistance mechanisms, including secondary and tertiary mutations, alternative pathway activation, *cMET* amplification, *HER2* amplification, EGFR nuclear trans-localization, oncogenic lncRNA regulation, and tumor microenvironment alterations^3, 8, 25–27^. The underlying mechanisms of EGFR-TKI resistance are not fully understood, mainly due to lack of approaches for complete elimination of EGFR in cells. Although EGFR nuclear translocation is one of the resistance mechanisms, whether HER2/3/4 nuclear translocation contributes to TKI resistance remains largely unknown. Therefore, we aimed to characterize a panel of *EGFR* knockout NSCLC cell lines and further to investigate other HER family receptors upon EGFR loss or acquired resistance. We show that cells with mutant *EGFR* behave differently than wild-type cells after *EGFR* disruption. We demonstrate that HER2 and HER3 overexpression and transcriptional activation of cyclin D1 are other resistance mechanisms to TKIs in NSCLC (Fig. 10).

**Fig.10.**
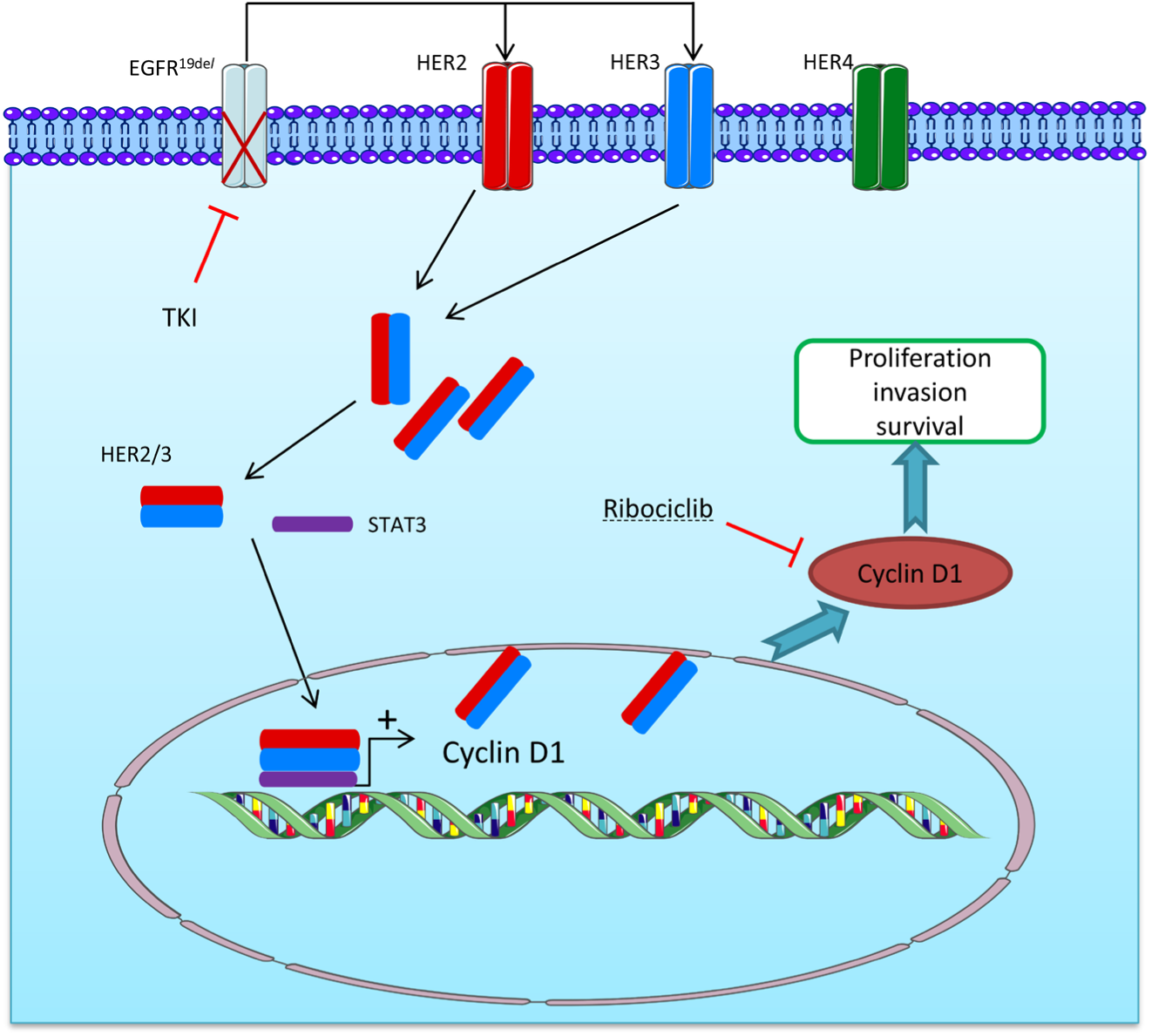
EGFR inhibition or disruption causes HER2/HER3 overexpression and nuclear translocation. Subsequently, HER2, HER3 and pSTAT3 form a heterodimer and transcriptional activation of cyclin D1, which contributes to cell survival and EGFR tyrosine kinase inhibitor resistance in non-small cell lung carcinoma.

Our results show that wild-type EGFR ablation resulted in a significant inhibition of cell proliferation, migration and 3D spheroid formation, but EGFR^19del^ ablation increased those processes. Wildtype EGFR plays an essential role in cell growth by a ligand-dependent mechanism. Therefore, knockout of EGFR^wt/wt^ leading to the downregulation of those abilities is not a surprise. Although *EGFR* exon 19 deletion is an activating mutation that can promote cell growth through a ligand-independent activation of tyrosine kinase domain^28^, disruption of mutant *EGFR* in H1650 cell line promoted cell growth and migration abilities. We indeed observed that the acute inhibition of EGFR using TKIs in mutant cells resulted in a dramatic suppression of cell proliferation. However, *EGFR^19del^* cell lines with long-term exposure to elortinib (HCC827-ER and H1650-ER) developed an even stronger proliferative ability and aggressiveness. These data suggest that EGFR^19del^ knockout may simulate EGFR-TKI resistance. This is in agreement with the fact that drug resistance leads to tumor hyper-progression^29^. A recent study provided a proof-of-concept of therapeutic benefits of *EGFR* knockout in *EGFR*-mutant NSCLC^30^. However, our study brings some concerns whether it is safe to remove mutant *EGFR* from tumor cells.

We observed significant changes in the levels of pERK(1/2) and pAKT upon EGFR ablation, inhibition, or acquired resistance. Firstly, pAKT levels were higher in A549 EGFR^-/-^, H1650 EGFR^-/-^ and HCC827-ER than in parental cells. Our data suggest that activation of the PI3K/AKT pathway may be an important compensation mechanism for tumor cell survival after EGFR loss. In agreement with our findings, a recent study showed that Akt activation drives acquired resistance to EGFR-TKI^31^. However, our study further revealed that activated Akt induced by EGFR ablation can be further decreased by elortinib treatment. In addition, pERK level significantly increased only in A549 *EGFR^-/-^* cells upon elortinib treatment, which might be due to off-target effects of erlotinib.

Our data revealed that HER2 and HER3 were significantly overexpressed in *EGFR^19del^* ablated and ER cells, but downregulated in *EGFR^wt/wt^* disrupted cells. Dysregulation of HER family receptors has been reported in patients treated with EGFR-TKIs and monoclonal antibodies, but the underlying mechanisms are not fully understood^32^. We showed that upregulation of HER family receptors upon TKIs treatment or EGFR ablation only occurred in *EGFR*-mutant tumor cells, which might be of importance in a clinical setting. Moreover, our findings, to our knowledge for the first time, further revealed a significant nuclear overexpression of HER2 and HER3 in NSCLC *EGFR^19del^* ablated or ER cells. Due to lack of an efficient approach for targeting nuclear protein, these findings may partially explain ineffectiveness of combination therapies by anti-EGFR and other HER family receptors.

We found a new mechanism of tumor cell survival through HER2/HER3/STAT3 complex binding and transcriptional activation of cyclin D1 in EGFR targeting in NSCLC (Fig. 10). Although cyclin D1 overexpression mediated by cyclinD1/CDK4/6 signaling and HER2/HER3/STAT3 nuclear translocation has been identified as an important resistance mechanism and therapeutic target in breast cancers^21, 33–35^, whether they also occur in lung cancer was unknown. Firstly, we showed that HER2 and HER3 have nuclear overexpression in mutant *EGFR* knockout and ER cell lines, which may compensate for EGFR loss. Secondly, we observed that HER2, HER3 and pSTAT3 form more heterotrimers in nuclei upon mutant *EGFR* ablation and erlotinib resistance. Thirdly, HER2/3/pSTAT3 complexes were enriched and bound to cyclin D1 promoter and subsequently activated cyclin D1 expression only after mutant *EGFR* knockout and erlotinib resistance. This resulted in an increased in S and G2/M phase cell population. However, we observed an opposite result in cells with wild-type EGFR. Furthermore, ribociclib was able to resensitize cells to erlotinib. Thus, our findings demonstrate that cyclin D1 overexpression mediated by HER2/3 nuclear translocation is a mechanism of TKI resistance in NSCLC. In addition, cyclin D1 high overexpression is associated with shorter overall survival and disease-free survival. Cyclin D1 might be a promising therapeutic target for patients with resistant NSCLC. Concomitant inhibition of EGFR and cyclin D1 may result in more prominent outcomes.

## Conclusions

In conclusion, our results show a distinct expression pattern of HER family receptors after EGFR loss in different NSCLC cells and it appears to be a mechanism of resistance to TKIs in NSCLC. Moreover, inhibition of wild-type and mutant EGFR showed distinguished impacts on other HER family receptors as well as tumor survival. In addition, cyclin D1 may be a potential therapeutic target for patients with resistant NSCLC.

## Materials and methods

### Cell culture and establishment of *EGFR* knockout (*EGFR^-/-^*) cell lines by CRISPR/Cas9

Four lung cancer cell lines (A549, H1299, H1650 and HCC827) were used in this study. A549 (*EGFR^wt/wt^*) was purchased from ATCC. H1299 (*EGFR^wt/wt^*) and H1650 (EGFR^19del^) were gifts from Dr. Klaas Kok (Department of Genetics, University Medical Center Groningen). The HCC827 (*EGFR^19del^)* cell line was kindly provided by Dr. Martin Pool (Department of Medical Oncology, University Medical Center Groningen). The *EGFR* knockout cell lines (A549 *EGFR^-/-^*, H1299 *EGFR^-/-^* and H1650 *EGFR^-/-^*) were generated by CRIPR/Cas9, as described^16^. All cell lines were cultured in Roswell Park Memorial Institute (RPMI)-1640 (Gibco, Carlsbad, USA) containing 1% (v/v) penicillin G/streptomycin (Invitrogen, Carlsbad, USA) supplemented with 10% (v/v) fetal bovine serum (FBS) (Thermo Fisher Scientific, Waltham, USA) in a humidified incubator at 37°C with 5% CO_2_. All human cell lines have been authenticated using STR (or SNP) profiling within the last three years. All experiments were performed with mycoplasma-free cells.

### Anti-tumor agents

Erlotinib and afatinib were purchased from Selleckchem (Munich, Germany). Ribociclib was purchased from LC laboratories (Woburn, USA). All drugs were dissolved in dimethyl sulfoxide (DMSO) and stored at −20°C.

### Establishment of EGFR-TKI resistant cell lines

EGFR-TKI resistant cell lines were established by long-term exposure to erlotinib (3 months). To set up EGFR TKI-resistant cell lines, HCC827 *EGFR^19del^* and H1650 *EGFR^19del^* were treated with erlotinib starting at 50 nM, followed by a stepwise dosage increase every 5 days up to 20 μM and 50 μM, respectively. The erlotinib-resistant (ER) cell lines HCC827ER and H1650ER were maintained in 2 μM erlotinib for long-term culture.

### Cell viability

Cell viability was measured using the MTS assay. Cells were seeded in flat bottom 96-well plates at a density of 1×10^3^ cells per well. After 24 hr, cells were treated with appropriate drugs at different doses for another 72 hr. Subsequently, CellTiter 96 Aqueous One Solution reagent (Promega, Madison, USA) was added to each well, according to manufacturer’s instruction. Plates were incubated at 37°C for 1hr. The optical density (OD) was determined at 490 mm wavelength using a Synergy H1 plate reader (BioTek, Winooski, USA).

### Cell proliferation

The colony formation assay was used to determine cell proliferation. Cells were seeded in 6-well plates at a density of 1×10^4^ cells per well and were allowed to grow for 4-5 days. Subsequently, cells were fixed with 4% (v/v) paraformaldehyde for 20 min at room temperature (RT) in the dark. After washing with phosphate-buffered saline (PBS), cells were stained with 0.5% (w/v) crystal violet for 15 min. Then, cells were washed three times with demineralized water and plates were dried at room temperature. Crystal violet was eluted by 10% (v/v) acetic acid and OD was measured at 590 mm wavelength using a Synergy H1 plate reader.

### Cell migration

To measure lateral migration, cells were seeded in a 6-well plate at a density of 7×10^5^ cell per well. After 24 hr incubation, a wound was gently scraped using a sterile 200 μl pipette tip. The detached cells were washed away with PBS. Pictures were taken at 0 hr, 12 hr, 24 hr, 48 hr using a CK2 inverted microscope (Olympus, Tokyo, Japan). The pictures were analyzed by ImageJ (National Institute of Health, USA).

To measure vertical migration, cells were seeded in the top trans-well inserts at a density of 1×10^4^ cells per insert. The inserts were gently put onto 24-well plates filled with 750 μl cell culture medium. Culture medium and remaining cells in the inserts were removed after 24 hr incubation at 37°C with 5% CO_2_. The insert membrane was fixed with 4% (v/v) paraformaldehyde and stained with 0.5% (w/v) crystal violet. Pictures of migrated cells were randomly taken using a CK2 inverted microscope. Subsequently, crystal violet was eluted with 10% (v/v) acetic acid and the OD590 value was measured using a Synergy H1 plate reader.

### Generation of a 3D-spheroid tumor model

To mimic actual tumor microenvironment, a 3D-spheroid tumor model was constructed, which has been detailed previously^17^. The U-bottom ultra-low attachment culture plate was used for the generation of a 3D-spheroid tumor model. Approximately 2×10^3^ cells per well were seeded into U-bottom ultra-low attachment 96-well plates. Plates were centrifuged at 300 g for 5 min to form a preliminary 3D spheroid tumor. After 5 days of incubation, 3D spheroid tumors were generated. The pictures were taken under the CKX41 microscope (Olympus, Tokyo, Japan) controlled by Leica Application Suite software. The image data were analyzed by ImageJ.

### RNA isolation, cDNA synthesis and qRT-PCR

Cells were harvested by trypsinization and RNA was isolated using Maxwell LEV simply RNA Cells/Tissue Kit (Promega, Madison, USA), according to the company’s protocol. A total of 500 ng RNA was reverse-transcribed to cDNA using the Reverse Transcription Kit (Promega, Madison, USA), according to the manufacturer’s instruction. The SensiMix SYBR Kit (Bioline, Taunton, USA) was used for qRT-PCR in a QuantStudio 7 Flex Real-Time PCR System (Thermo Fisher Scientific, Waltham, USA). Data were analyzed by QuantStudio Software V1.3 (Thermo Fisher Scientific, Waltham, USA). α-tubulin mRNA level was determined as an internal reference for data normalization.

### Western blot analysis

Cells were harvested by trypsinization and lysed on ice in RIPA buffer containing 1x PhosSTOP and protease inhibitor (PI) cocktail (Roche, Mannheim, Germany) for 20 min. The lysates were sonicated for 10-20 seconds to reduce the viscosity. Next, samples were centrifuged at 16,000 RCF, 4°C for 20 min to remove insoluble residues. Protein concentration was determined using the BCA Protein Assay Kit (Pierce, Rockford IL, USA), according to the manufacturer’s instruction. Thirty micrograms denatured protein of each sample was mixed with SDS buffer and loaded onto a pre-cast 4-12% NuPAGE Bis-Tris gel (Invitrogen, USA). Later, proteins were transferred to a polyvinylidene difluoride (PVDF) membrane. After blocking with 5% (w/v) skimmed milk in PBST (Phosphate Buffered Saline with Tween 20) at room temperature for 1 hr, the membrane was incubated overnight at 4°C with appropriate primary antibody. Then, the membrane was treated with HRP-conjugated secondary antibody for 1 hr at RT. After each step the membrane was washed three times with PBST. Blots were lighted up with enhanced chemiluminescence (ECL) solution (GE Healthcare, Amersham, UK). Pictures were captured using a chemi genius II bio-imaging system.

### Nuclear isolation and co-immunoprecipitation (co-IP)

The nuclear isolation and co-immunoprecipitation (co-IP) assay has been detailed elsewhere^18, 19^. Briefly, 3×10^8^ Cells were harvested under non-denaturing conditions for nuclear isolation, followed by washing with PBS. Cells were resuspended in 400 μl hypotonic lysis buffer and incubated on ice for 20 min. Nuclei were pelleted by centrifugation at 500 g, 4°C for 10 min. Supernatant contained the non-nuclear fraction and the pellet was the nuclear fraction.

For co-IP, samples were gently washed three times with clod PBS containing 0.1% CA-630 and 1x protease inhibitor (PI) to minimize cytoplasmic contamination. Nuclear protein samples were extracted using 300 μl RIPA buffer containing 1x PhosSTOP and PI. After 20 min incubation on ice, samples were sonicated and centrifuged at 16,000 RCF, 4°C for 20 min. The concentration of non-nuclear and nuclear protein samples was determined by BCA Protein Assay Kit, according to the manufacturer’s instruction. Approximately 300 μg of cell lysate at 1-1.5 mg/ml was then pre-cleaned by rotation incubation with 20 μl 50% Protein A/G PLUS-Agarose (Santa Cruz Biotechnology, Texas, USA) at 4°C for 60 min. Subsequently, appropriate antibody was added to pre-cleaned samples, followed by overnight rotation incubation at 4°C. Next, 30 ul 50% Protein A/G PLUS-Agarose was added to each sample. The mixtures were incubated at 4°C for overnight with rotation. The pellets were collected by centrifugation at 1000 g for 3 min at 4°C and washed five times with 500 μl RIPA buffer. The pellets were well mixed with 20 μl 4x SDS buffer and denatured by heating at 95°C for 10 min. Denatured protein samples were analyzed by western blot.

### Sequential chromatin immunoprecipitation (sequential ChIP)

The sequential ChIP-qPCR has been described elsewhere^20–22^. Cells were grown in a T175 cell culture flask to 90% confluency. Freshly prepared 11% (v/v) formaldehyde solution was directly added to the flask at a 1% formaldehyde concentration. The flasks were gently swirled at room temperature for 20 mins for protein-chromatin cross-link formation, followed by addition of 2.5 M glycine to each flask to reach a 0.1 M final concentration. After washing with PBS, cells were harvested under non-denaturing conditions. To isolate the nuclei, the cell pellet was resuspended in 900 μl lysis buffer and lysed by pipetting. Then, cell lysates were washed once with 1 ml ChIP lysis buffer. Chromatin fragments (100-1000 bp) were obtained using micrococcal nuclease digestion followed by gel electrophoresis. Thirty percent of the chromatin solution (unbound sample) was stored at −20°C and the remaining 70% of chromatin solution was incubated with appropriate primary antibody overnight at 4°C with rotation. Then, 30 ul of 50% Protein A/G PLUS-Agarose was added. After a 3 hr incubation at 4°C with rotation, agarose beads were collected and sequentially washed three times with 1 ml RIPA buffer and twice with 1 ml TE buffer. The immune complex was eluted with 10 mM dithiothreitol (DTT) in 75 μl TE buffer at 37°C for 30 mins. The eluted immunoprecipitate was diluted 20 times with dilution buffer. The second round ChIP was performed as the first round. The re-immunoprecipitation was eluted and cross-links were removed with elution buffer overnight at 65°C. The de-cross-linked chromatin and the unbound sample were sequentially incubated with RNase A (10 mg/ml) for 1 hr at 37°C, and proteinase K (20mg/ml) for 2 hr at 56°C. Finally, DNA fragments were purified with QIAquick® PCR purification kit (Qiagen, Hilden, Germany) according to the manufacturer’s instruction. Primers are listed in Supplementary Table S1. The qPCR amplifications have been described elsewhere^22^.

### Kaplan-Meier survival and correlation analysis of patient data

Data for survival analysis was retrieved from The Cancer Genome Atlas (TCGA) using a standard processing pipeline. Cumulative disease free survival of NSCLC (LUAD, LUSC) was analyzed using the Kaplan-Meier method with log-rank test. Correlation between the expressions of different genes was calculated using Spearman correlation (non-log scale) based on Gene Expression Profiling Interactive Analysis (GEPIA)^23^. A two-sided p-value less than 0.05 was considered statistically significant.

### Data analysis

Data were derived from three independent experiments and were presented as mean ± SD. P-values were determined by GraphPad Prism software v.6.0.1 using one-way ANOVA with multiple comparisons and Tukey’s correction. * P≤0.005, *** p≤0.0005, **** p≤0.0001.

## Abbreviations

Co-IP: Co-immunoprecipitation
CHIP: Chromatin immunoprecipitation
CRISPR: Clustered Regularly Interspaced Short Palindromic Repeats
ER: Elortinib resistance
EGFR: Epidermal growth factor receptor
*EGFR*^19del^: Epidermal growth factor receptor exon 19 deletion
*EGFR^-/-^*: EGFR knockout
NSCLC: Non-small cell lung carcinoma
OS: overall survival
DFS: disease-free survival
pEGFR: phosphorylated epidermal growth factor receptor
STAT3: Signal transducer and activator of transcription 3
TCGA: The Cancer Genome Atlas
TKI: Tyrosine kinase inhibitor

## Declarations

### Ethics approval and consent to participate

Not applicable.

### Data availability

The data that support the findings of this study are available from the corresponding author upon reasonable request.

### Consent for publication

Not applicable.

### Authors’ contributions

H.J.H. and B.L. designed the study. B.L. and D.C. performed molecular and cellular experiments. S.P.C analysed and interpreted the patient data. A.S. and B.L. established EGFR knockout cell lines. D.C. managed and analysed the figures and data under the supervision of B.L and H.J.H. B.L. wrote the manuscript. All authors read and approved the final manuscript. H.J.H supervised this project.

## Acknowledgements

We thank Petra E. van der Wouden and Rita Setroikromo for technical support. We thank Martijn Zwinderman for suggestions of ChIP-qPCR study. Bin Liu and Shipeng Chen have received PhD research fellowship from the China Scholarship Council.

---

**Supplementary Table S1.**
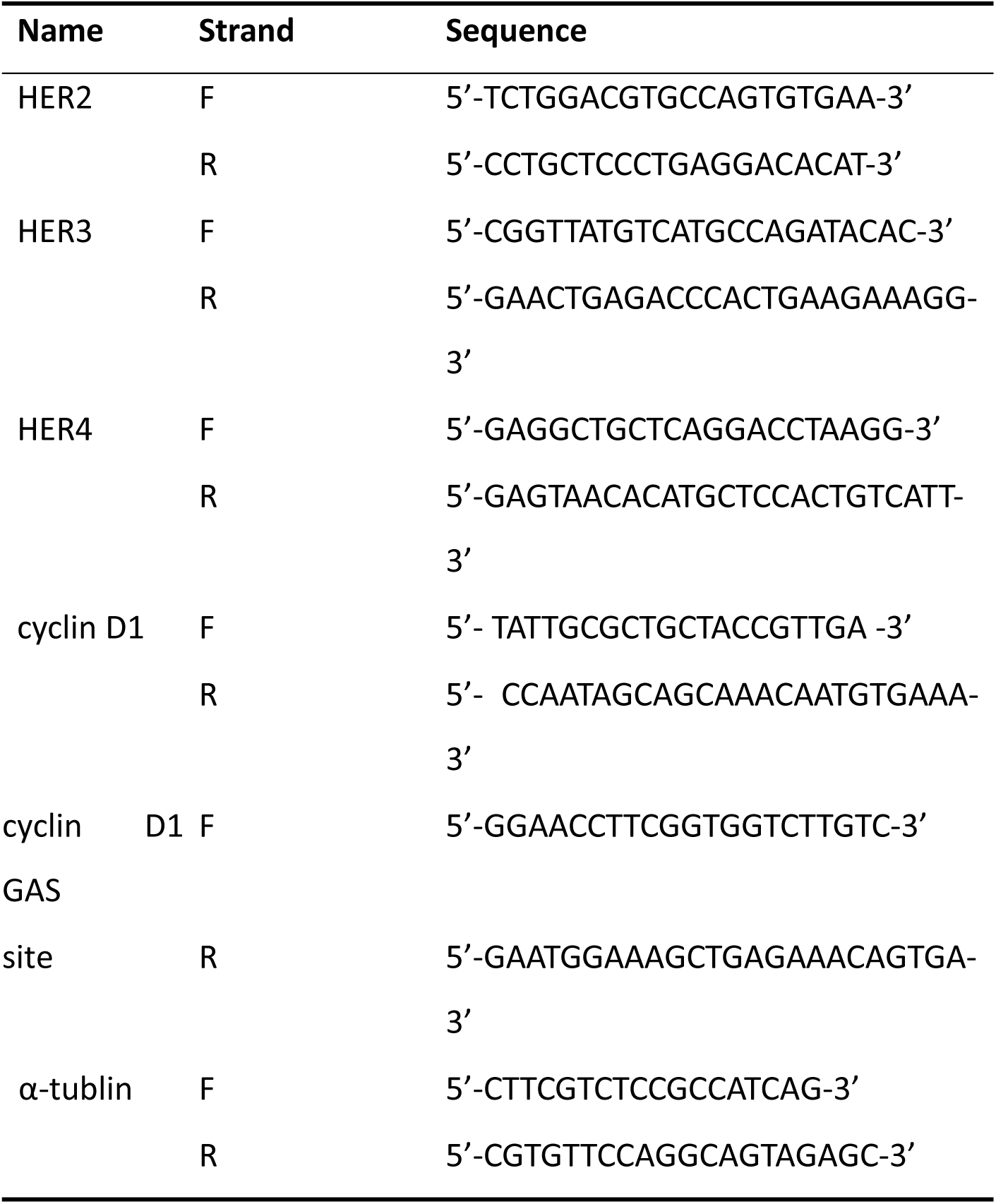
List of the primer sets used for qRT-PCR and CHIP-qPCR.

**Supplementary Fig. S1.**
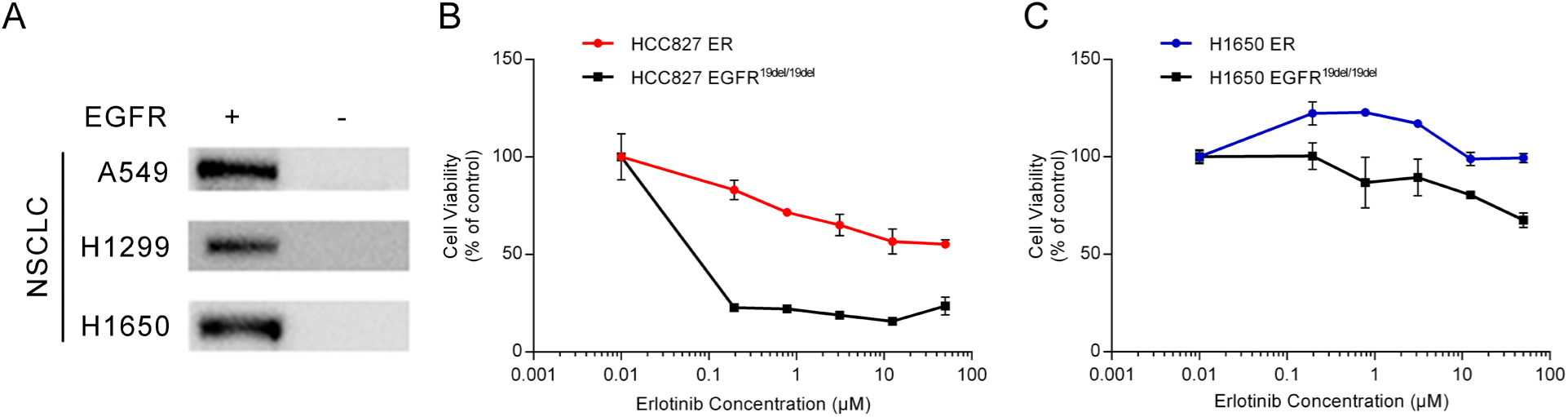
Establishment of *EGFR* knockout and erlotinib-resistant NSCLC cell lines. **A.** Western blot analysis of *EGFR* expression in in *EGFR* knockout and original cells. **B.** Cell viability analysis of H1650, H1650ER, HCC827 and HCC827ER cell lines 72 hours after erlotinib treatment.

